# SLEAP: Multi-animal pose tracking

**DOI:** 10.1101/2020.08.31.276246

**Authors:** Talmo D. Pereira, Nathaniel Tabris, Junyu Li, Shruthi Ravindranath, Eleni S. Papadoyannis, Z. Yan Wang, David M. Turner, Grace McKenzie-Smith, Sarah D. Kocher, Annegret L. Falkner, Joshua W. Shaevitz, Mala Murthy

## Abstract

The desire to understand how the brain generates and patterns behavior has driven rapid methodological innovation to quantify and model natural animal behavior. This has led to important advances in deep learning-based markerless pose estimation that have been enabled in part by the success of deep learning for computer vision applications. Here we present SLEAP (Social LEAP Estimates Animal Poses), a framework for multi-animal pose tracking via deep learning. This system is capable of simultaneously tracking any number of animals during social interactions and across a variety of experimental conditions. SLEAP implements several complementary approaches for dealing with the problems inherent in moving from single-to multi-animal pose tracking, including configurable neural network architectures, inference techniques, and tracking algorithms, enabling easy specialization and tuning for particular experimental conditions or performance requirements. We report results on multiple datasets of socially interacting animals (flies, bees, and mice) and describe how dataset-specific properties can be leveraged to determine the best configuration of SLEAP models. Using a high accuracy model (<2.8 px error on 95% of points), we were able to track two animals from full size 1024 × 1024 pixel frames at up to 320 FPS. The SLEAP framework comes with a sophisticated graphical user interface, multi-platform support, Colab-based GPU-free training and inference, and complete tutorials available, in addition to the datasets, at sleap.ai.

## 1. Introduction

Quantitative measurements of animal motion are foundational to the study of animal behavior [1, 6, 11]. Methods for *pose estimation*, the task of predicting the location of anatomical landmarks in images, have rapidly grown in popularity as a state-of-the-art technique for behavioral quantification across disciplines including neuroscience [24] and ecology [10]. Although adaptations of deep learning-based approaches originally developed for human pose estimation have made animal pose estimation possible [22, 26, 17], reliably tracking multiple, interacting animals and their pose remains a challenging problem, presenting an impediment to studies of social behaviors.

The generalization of the pose estimation task to the domain of multiple individuals (i.e., **instances**) can be broken down into three distinct sub-tasks:

i. **Landmark localization**: The retrieval of coordinates of each landmark from the image. In the multi-instance setting, there may be more than one detection of each landmark type (e.g., multiple necks). Localization is more challenging in the context of socially behaving animals due to increased occlusions from close interactions, along with imaging constraints such as low resolution and contrast (Figure 1a-b).
ii. **Part grouping**: The grouping of detected landmarks into distinct sets associated to each individual. This requires more information than simply the location of the landmarks, resulting in the *part grouping problem* (Figure 1c). In the context of socially behaving animals, intersecting or overlapping parts increase the difficulty of the grouping problem (Figure 1a-b).
iii. **Temporal association**: The association of grouped landmark sets across video frames such that landmarks belonging to each individual are consistently assigned the same identity. The *temporal association problem* requires defining a metric of affinity between instances across frames and matching them over time (Figure 1d). This is also known as *pose tracking* and is similar to multi-object tracking with the additional constraint of having multiple point types (landmarks) that form distinct sets.

**Figure 1:**
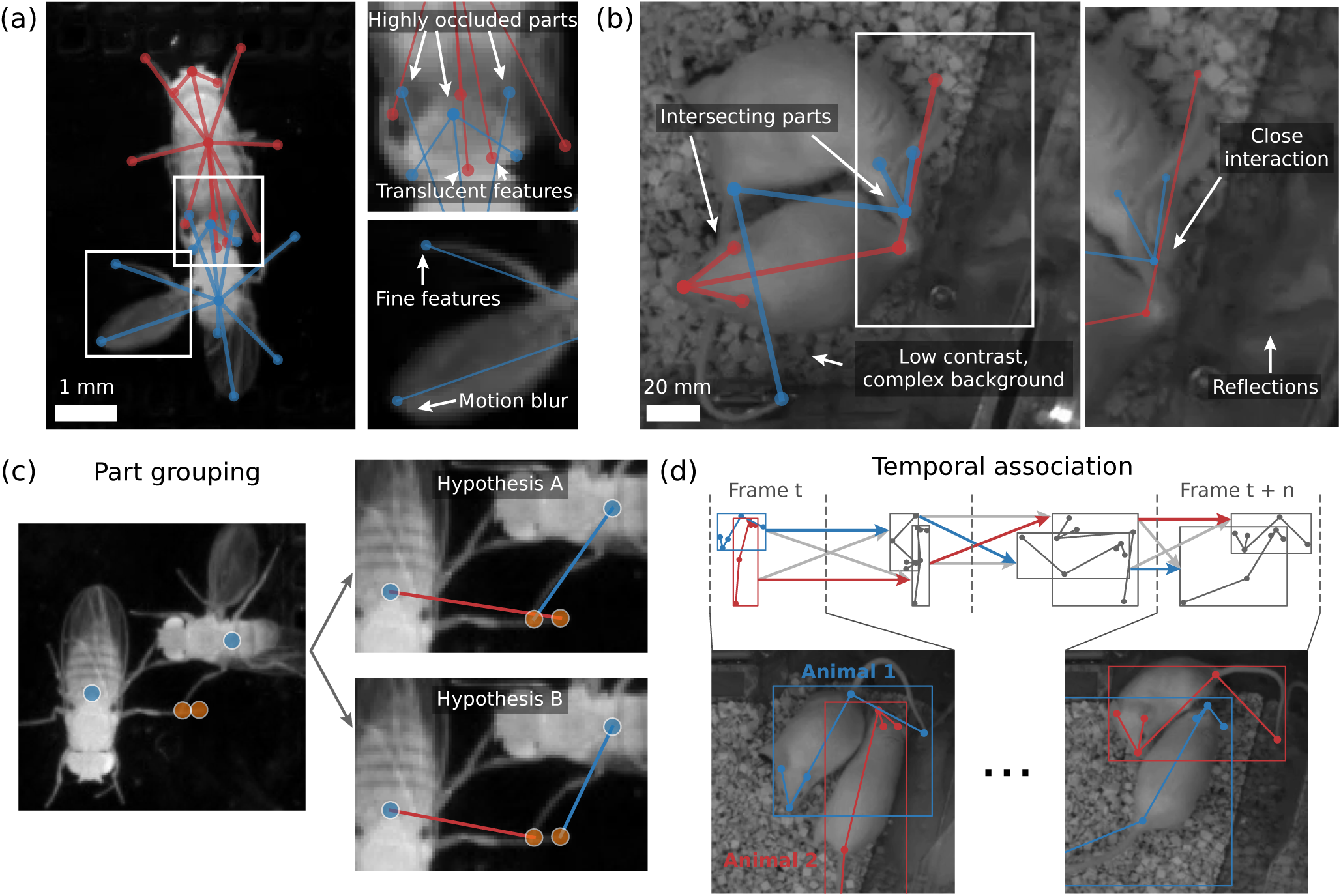
Unique challenges in pose tracking of socially behaving animals. **(a)** Fruit flies (*D. melanogaster*) engaged in a courtship interaction. Part detection during social behaviors such as tapping is difficult due to frequent occlusion, including through partially translucent body parts such as wings (*inset, top*). Even at high spatial resolution (∼30px/mm), anatomical features such as the leg tips occupy ∼3px, requiring the use of image features at the finest resolution, while their fast movements introduce motion blur even at high temporal resolution (150 FPS), making it difficult to precisely localize body parts (*inset, bottom*). **(b)** Mice (*M. musculus*) engaged in anogenital sniffing, a common social recognition behavior. The flexible morphology of these animals leads to frequent intersections of body segments, particularly the tail, making part association considerably more challenging. Naturalistic behavior is typically recorded in infrared and in a home cage environment which results in a low contrast, heterogeneous and dynamic background due to the bedding material. During social behavior, mice interact at close ranges which results in frequent occlusions; additionally, they spend most of their time near the walls which results in reflections that must be distinguished from real part detections (*inset, right*). **(c)** The part grouping problem emerges out of the generalization of pose estimation to multiple instances. When there are multiple detections of the same body part, such as the thorax (*purple*) or leg (*orange*), grouping these such that each distinct subset of body parts belongs to the same animal requires resolving competing hypotheses of how body parts may be connected (*right*). **(d)** The temporal association problem is the time-dimension analogue of the part grouping problem. Given frame-wise instance groupings of pose detections, consistent identities must be assigned to detections of the same animals across frames. Matching instances across frames requires solving an assignment problem that is robust to frequent crossings of both individual landmarks as well as the bounding box (*top*, candidate assignments in gray, correct assignments in colors). Animals may adopt very similar poses only a few frames apart (*bottom*), making it difficult to find globally optimal associations.

In this work we present a method to address the problems of *multi-animal pose tracking* and describe an implementation of our approach within a general-purpose software framework we term **SLEAP** (**S**ocial **L**EAP **E**stimates **A**nimal **P**oses).

The main contributions of our work are as follows:

1. We develop a software framework with a sophisticated GUI-driven workflow for labeling, training, tracking and proofreading social behavioral datasets.
2. We describe the two major classes of multi-instance pose estimation approaches (top-down and bottom-up) and their generalization to the unconstrained animal domain.
3. We describe an algorithm for pose tracking that can be configured to solve the temporal association problem with both image- and motion-based cues.
4. We provide a principled adaptation of the most commonly employed human pose estimation and tracking accuracy metrics to the animal domain.
5. We describe a set of high level hyperparameters that can induce specialized neural network architectures to meet the requirements of a given dataset.
6. We demonstrate the performance of neural network architectural hyperparameters and multi-instance approaches through extensive experiments across multiple datasets representing diverse social animals, to guide practitioners in training models on their own data.
7. We explore the impact of transfer learning on multianimal pose estimation performance.
8. We explore the speed-accuracy trade-off of different models and approaches, attaining maximum end-to-end multi-animal inference speeds of 430 FPS on a single GPU.

The open-source software framework implementing these methodological advances, as well as labeled datasets and trained models, are freely available at sleap.ai.

## 2. Related work

### 2.1. Human pose estimation and tracking

Previous work in multi-human pose estimation has inspired some of the approaches we employ here. The bottom-up multi-instance pose estimation technique we employ is based off of the part affinity fields representation and matching algorithm employed in the widely used human pose estimation framework OpenPose[7]. Our top-down approach and part of our tracking algorithm are inspired by the idea of flow shifting described in Xiao et al. [33]. Many of the metrics we use to evaluate both our multi-instance pose estimation accuracy as well as tracking come from the standards put forth in the PoseTrack benchmark [2].

### 2.2. Animal pose estimation

Early work on animal pose estimation extended deep learning algorithms designed for human subjects for use on animals with different body plans and morphologies. These include DeepLabCut[22], LEAP[26], and DeepPoseKit[17], but none were explicitly designed for use with multiple animals.

Methods for multi-animal pose estimation can be split into two categories: *top-down*, where animal instances are first detected and isolated before finding their individual body parts; and *bottom-up*, where body parts are first detected and then grouped into instances. DeepLabCut, LEAP, and DeepPoseKit, have all been adapted to perform multi-animal pose estimation in a top-down framework, but the process of identifying animal instances must be performed separately. More recent work using 3D data also employed top-down approaches for multi-animal pose but only in single species or specialized experimental conditions [5, 15, 12]. Other work has focused on bottom-up approaches using rodents, but these methods have not been shown to generalize to other types of animals [3, 19].

SLEAP is a framework that is designed for general-purpose 2D multi-animal pose estimation and tracking. We employ both top-down and bottom-up frameworks and conduct systematic experiments to compare these approaches and their effectiveness with a diversity of behavioral datasets from multiple species. Importantly, we find that depending on the dataset, either top-down or bottom-up methods give superior performance.

## 3. Method

### 3.1. Framework

The SLEAP multi-animal pose tracking framework is composed of a series of steps that form a standard workflow starting from data input and resulting in trained pose estimation models and pose tracked videos. The typical use workflow will sequentially step through each of the modules in the framework (Figure 2a, from left to right):

i. **Data input**. Unlabeled and unprocessed videos, e.g., multiple sessions from a experimental setup. Videos can be loaded from most video formats including MP4, AVI, image folders, HDF5, or imported from common project formats such as DeepLabCut [22], Deep-PoseKit [17] or standardized pose estimation dataset formats like MS-COCO [21].
ii. **Interactive labeling**. The user creates a new project in the cross-platform desktop labeling GUI, containing the videos, labels, and the specification of which landmarks to track. Labeling is performed by dragging landmark markers onto their corresponding positions for each animal in the image and does not require a GPU or a high performance machine. Each animal in an image is labeled with a distinct set of landmarks — the user does not need to keep track of animal identities at this stage. SLEAP can intelligently “suggest”” frames to label by analyzing their image content to maximize sample diversity [26].
iii. **Neural network model training**. Once as few as ∼10 frames have been labeled, predefined or user-specified neural network configurations can be used to train the initial neural network for multi-animal pose estimation. Users can either select from a list of pre-made default configurations that are designed for the most common use case, or can be guided through the configuration process, with documented descriptions of the hyperparameters accessible within the GUI, along with visualizations and previews of the outputs that will be produced by a given configuration. If the user is not labeling on a machine equipped with a GPU, SLEAP can export the project file packaged with the image data corresponding to the labels into a single file that can be uploaded to an institutional compute cluster, remote cloud instance, or even Google Colab which provides free GPU access. The user can install SLEAP remotely with a simple `pip install sleap` command, train the neural network by uploading the desired configuration, monitor the training via TensorBoard, and then download the resulting trained model. These steps can be done via an interactive Python notebook or via commandline interface (CLI) for batch scheduling. On machines with local GPU access, training can be done interactively and training progress monitored directly within the SLEAP GUI, including visualizations of predictions during model training. Training speed depends on the dataset and network configuration, but predefined templates will typically converge between 10 and 30 minutes for initial training. After training, the resulting model folder will contain the neural network weights, training logs, visualizations, and accuracy metrics, as well as a cached copy of the labels used to train the network and the training configuration, so that the training procedure that generated a saved SLEAP model is fully reproducible.
iv. **Pose estimation**. Once the network is trained, new poses can be predicted for all animals on a per-frame basis. Predictions can optionally be automatically generated on suggested frames or on user-selected frames. Prediction results after the initial round of training will likely be misplaced, swapped, or missed entirely, however correcting these inaccurate predictions will take *considerably* less time than labeling images from scratch. New predictions can be generated on single frames or entire videos or datasets, depending on user specification. Pose predictions are saved in the project file for future labeling sessions. Continued labeling can further take advantage of the outputs produced by trained models by sorting predictions by their confidence scores, where low confidence predictions may indicate deficiencies in the labels, for example, frames where the animals adopt poses not included in the training data. After adding more prediction-assisted labels to the dataset, the user can repeat the training procedure to train a more refined model with the added labels. The new model will then be able to generate predictions that are more accurate and require less time to correct, rapidly speeding up the process of labeling a new dataset. This is called **human-in-the-loop** training and forms the basis of expedient SLEAP workflows.
v. **Tracking**. Once the user is satisfied with the accuracy of the pose estimation model, SLEAP can run a separate tracking algorithm designed to operate on frame-wise predictions to associate predictions across frames. This tracker does not require training and can take advantage of both image data and motion cues to minimize tracking errors. Further post-processing utilities can improve the rate of tracking errors if the user is able to provide additional constraints such as the maximum number of animals or frame regions to ignore. This tracker can be run interactively on-demand as a subsequent step to the pose estimation pipeline, or as part of a batch processing workflow via the CLI or the Python API.
vi. **Proofreading**. After tracking is performed, predictions for new videos not in the labeling project are saved to their own SLEAP labels file. This file can be opened by the SLEAP GUI to inspect the predicted tracks and correct tracking or pose estimation mistakes if there are any. Tracking errors that may require proofreading typically track swaps due to prolonged close interactions between animals with occlusion or splits in tracks due to the animal being out of the frame or occluded for an extended period of time. SLEAP can compute a variety of different metrics that can help to spot tracking problems in longer videos, such as body part velocities (peaks may indicate track swaps) or low prediction confidence (indicating persistent animal occlusions).

**Figure 2:**
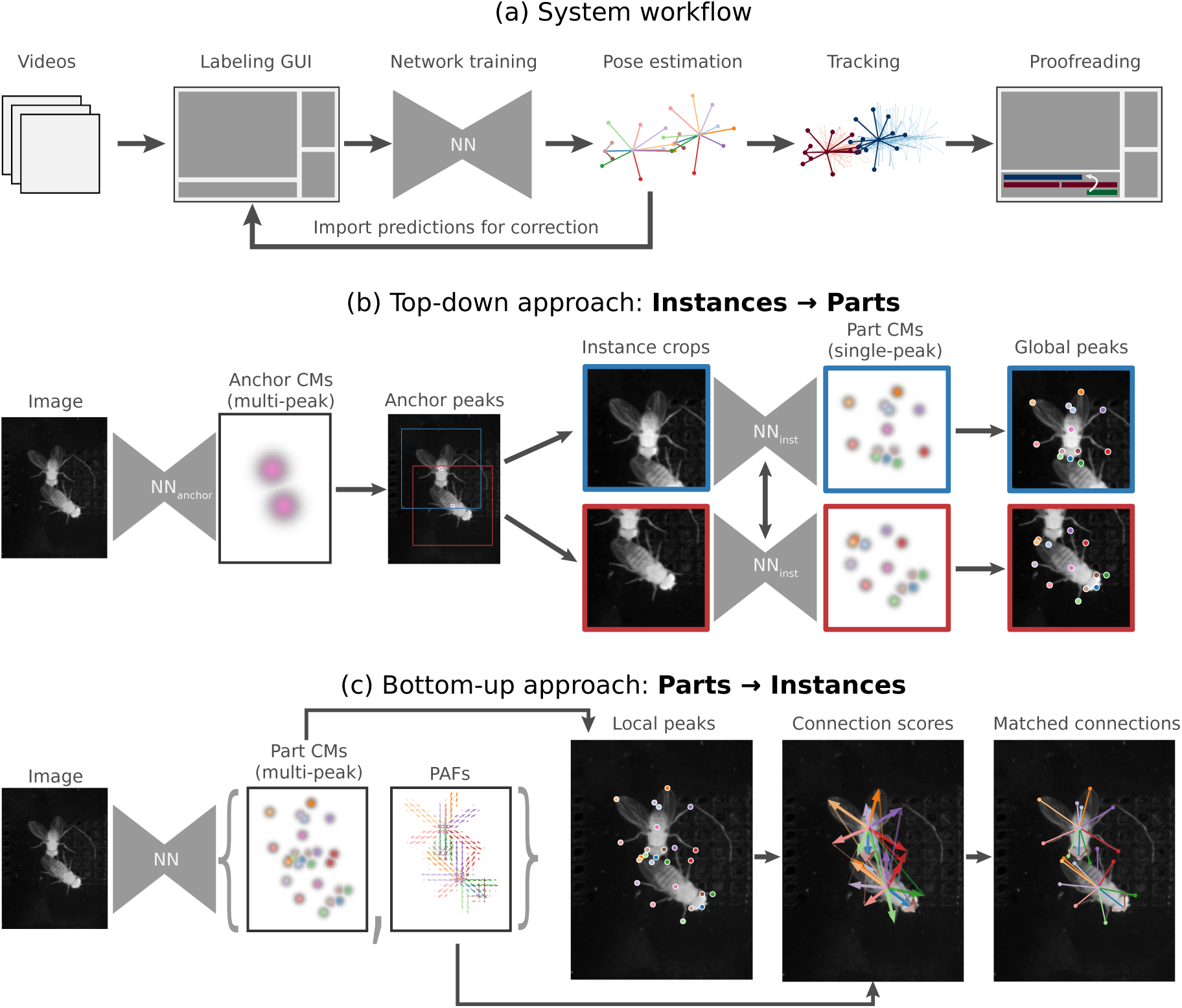
Overview of the SLEAP framework. **(a)** The standard workflow receives raw videos as inputs and produces multi-animal pose tracks as outputs. **(b)** The top-down approach to multi-instance pose estimation first finds instances and crops them from the full frame, then detects the individual’s body parts within each crop. In the first stage, a network (*NN*_*anchor*_) finds the instances by predicting multi-peak confidence maps of anchor points. The second stage network (*NN*_*inst*_) predicts single-peak confidence maps for the center instance of each crop. **(c)** The bottom-up approach finds all parts within the full image, then uses a connectivity metric to group them into instances. Both multi-peak confidence maps and part affinity fields (PAFs) are predicted by a single network (*NN*).

### 3.2. Landmark localization

The position of each landmark from the labeled data is encoded for network training by a 2D array that we refer to as a **part confidence map** (CM). For each body part coordinate **x**_*i*_ ∈ ℝ^2^, the value of the confidence map at pixel **x**_*p*_ ∈ ℝ^2^ is given by an unnormalized 2D Gaussian distribution,

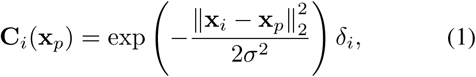

where *σ* is a fixed scalar controlling the symmetric spread of the distribution, and *δ*_*i*_ is equal to 0 when the body part is labeled as *“not visible”* and 1 otherwise.

The confidence maps are evaluated at each image grid pixel coordinate **x**_*p*_ ∈ {((*x, y*) : *x* ∈ {0, …, *W*}, *y* ∈ {0, …, *H*}} where *W* and *H* are the image width and height, respectively. The grid can be subsampled to generate lower resolution confidence maps as targets for neural networks, trading off spatial resolution for decreased memory usage and compute cost. For an animal with *J* body part types (e.g., head, thorax, etc.), we generate *N* confidence maps which are stacked along the channels axis such that the full confidence map’s tensor **C** is of shape (*H/s*_*o*_, *W/s*_*o*_, *N*), where *s*_*o*_ is the output stride of the network. Body parts that are marked as *“not visible”* during labeling are represented by a confidence map filled with zeros. We set *σ* = 1.0 and scale by the output stride to maintain a fixed scale with respect to the image resolution. For images with multiple instances of each body part type, the part confidence maps for each instance are combined by taking their maximum value at each pixel which helps to separate closely-spaced peaks [7].

The set of confidence maps from the labeled data is used to train the neural network which then predicts confidence maps for novel data. The confidence map representation has the benefit of enabling fully convolutional neural network architectures which are both efficient and easier to train than networks that directly regress the coordinates of each body part [32]. The trade-off is that the coordinates must be computed from the confidence maps at inference time (i.e., when the model is predicting new confidence maps).

For single-instance confidence maps, we decode the coordinates by finding the global peak, i.e., the coordinates of the confidence map pixel with the highest value. For multi-instance confidence maps, we employ local peak finding, where we define a pixel as being a local peak if it is greater than its 8 neighbors. In practice, we employ *non-maximum suppression* computed using a 2D grayscale dilation (maximum) filter with kernel

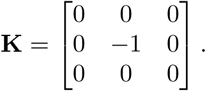

**K** is convolved with the confidence map, producing a tensor whose elements contain the maximum of each 3 × 3 patch excluding the central pixel. The pixels in the confidence map with higher values than those in the dilated maps are considered local peaks. In both global and local peak finding, we exclude peaks whose confidence map values fall below a fixed threshold which we set to 0.2 in order to retain somewhat low confidence predictions, but exclude points that are reliably predicted as *“not visible”*.

Since both of these peak finding methods can only yield peak coordinates at the resolution of the confidence map grid, localization accuracy is limited by the grid sampling interval. This quantization error is particularly problematic for models with larger output strides (i.e., lower resolution confidence maps), so we employ subpixel refinement to improve the peak coordinate localization. We leverage integral regression [25] to compute real-valued offsets by taking the weighted average of the local patch of the confidence maps around each grid-aligned peak.

### 3.3. Multi-instance approaches

To deal with the **part grouping problem** inherent in multi-animal pose estimation, we explore two types of approaches: *top-down* and *bottom-up* (Figure 2b-c). Both of these methods are widely employed in the human pose estimation literature and provide distinct trade-offs in terms of accuracy and performance. It has not been previously reported which may be better suited to the domain of multi-animal pose estimation, so we explored both.

#### 3.3.1 Top-down

In the top-down approach, the instances are first detected within the full resolution image and each instance is cropped. Each of the resulting crops will be centered on an instance, but may contain pixels that belong to other instances. This centering is crucial as it provides spatial context to the second stage of the top-down approach, serving as an indicator of which instance’s body parts to predict within the cropped image. In our framework, we select a labeled body part type to use as an *anchor*, ideally one close to the center of the animal’s bounding box and infrequently occluded (if occluded, the centroid of the bounding box of the remaining parts is used as the anchor). The anchored part serves as the target for the first stage neural network (*NN*_*anchor*_) which is trained to predict *multi-peak* confidence maps corresponding to the anchor part of all animals in the frame (Figure 2b, left). Typically, this network is trained on downsampled (0.25× or 0.5×) full frame images since coarsely locating the animals does not require high spatial resolution and saves on compute cost. Anchor part confidence maps are converted to coordinates using local peak finding and cropped from the full resolution images with a fixed bounding box size computed automatically from the labels.

Once the instance-anchored crops are produced by the first stage, the second stage essentially treats them as single-instance images. In this stage, we train a separate neural network (*NN*_*inst*_) that takes an instance-anchored image and predicts single-peak confidence maps **only for the anchored instance**. The confidence maps are converted into coordinates using global peak finding as only a single set of body parts are expected (Figure 2b, right). This network implicitly addresses the part grouping problem by leveraging the location of the body parts relative to the anchor part (i.e., the image center) as a cue for which body part to predict confidence maps for when multiple of the same body part type may be present within the crop.

This form of implicit modeling of the geometry between body parts is simple and has been employed in the animal pose literature previously [26, 17]. The disadvantages of the top-down approach are that it fails to capture global contextual information present in the relationship *between* instances, is limited by the accuracy of the first stage detector, and requires a full forward pass through the second stage network (*NN*_*inst*_) for each animal detected (though this may actually require *less* computation for images with few animals that occupy a small fraction of the image).

#### 3.3.2 Bottom-up

For the bottom-up approach, we employ an image-based representation of the connectivity between body parts that has been previously described for human pose estimation called *part affinity fields* (PAFs) [7]. This representation captures the relationship between body parts explicitly by encoding a vector field which locally points from each *source* body part to each *destination* body part. This vector field is stored as two 2D images, one for each component in the *x, y*-plane. In order to generate the PAFs from labeled data, the user must define a directed graph that connects all body parts to be tracked which we refer to as the *skeleton*. This skeleton graph must form a *spanning arborescence*, i.e., each node (body part) must have exactly one parent (except for a single root node), but may have multiple children. This is required for tractably solving the partitioning problem, which simplifies to a series of bipartite matching problems for arborescences. This is an important consideration when defining skeletons for new morphologies or datasets as body parts that are descendants of a missed body part will fail to be grouped correctly with the remaining instance parts. In practice, we attempt to create skeletons with the smallest depth to reduce dependency across sets of body parts.

A skeleton is defined as *S* = (*N, E*), where *N* is the set of *n* nodes (body parts) and *E* is the set of (*s, d*) tuples denoting the directed edge (connection) from a source node *s* ∈ {1, …, *n*} to a destination node *d* ∈ {1, …, *n*} \ {*s*}. The direction at each point in the PAF derived from labeled data is generated from the coordinates of the body parts in labeled images by computing the distance-weighted edge unit vector for each edge *e* at each image grid coordinate **x**_*p*_,

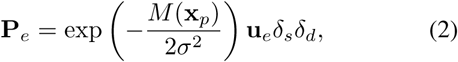

where **x**_*s*_ and **x**_*d*_ are the coordinates of the source and destination nodes, respectively. Similarly to confidence maps, **x**_*p*_ may come from a subsampled image grid, *σ* controls the spatial spread of the PAF, and *δ*_*s*_ and *δ*_*d*_ are the visibility flags for the source and destination nodes, respectively. The edge unit vector **u**_*e*_ is defined as the source-centered direction vector

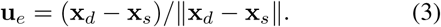

The magnitude, *M*, at each point in the PAF is defined as the Euclidean distance between the grid point **x**_*p*_ and its projection 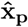 onto the line segment formed between **x**_*s*_ and 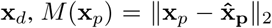, where 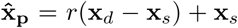 and

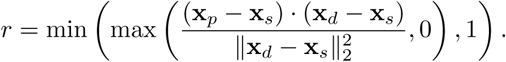

We note that the original description of PAFs [7] uses a hard threshold to compute the distance weighting, but we adopt a Gaussian instead as a means of scaling the relative contribution of pixels as a function of distance from the edge, resulting in smoother PAFs when animals are closely interacting.

PAFs computed for a given edge are combined for multiple instances by summation. After PAFs are generated for all edges in the skeleton, the full set of PAFs for the image **P** are of shape (*H/s*_*o*_, *W/s*_*o*_, 2 |*E*|), formed by concatenating all of the individual edge PAFs, which contain the *x-* and *y-* components of the vectors along the third axis.

In the bottom-up approach, a single neural network takes the full image as input and outputs both the PAFs and the multi-peak part confidence maps encoding the location of all body parts across all instances (Figure 2c, left). By predicting both of these representations, the network explicitly separates the task of localization and grouping, where for one representation it must only learn to predict “what” a body part is (CMs), whereas for the other it must learn the relationship between them (PAFs). This is in contrast to the top-down approach, where the relationship between body parts is implicitly encoded in the cropping.

After CMs are converted to peaks via local peak detection, sets of candidate source and destination peaks are grouped via greedy bipartite matching using the PAFs to compute the score of each putative connection (Figure 2c, right). For each pair of source and destination nodes, a line integral is computed by sampling 10 evenly spaced points between source and destination coordinates in the predicted PAFs. The score for the connection is calculated as the average dot product between the sampled vectors 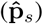 and the unit normalized vector formed between the predicted source 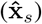 and destination points 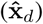 in the candidate connection,

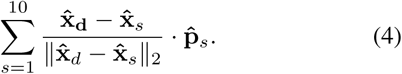

Once all pairs of connections are scored, instances are assembled by growing its skeleton edge by edge, assigning source candidates to destination candidates via Hungarian matching. The globally optimal matching is guaranteed through this local greedy procedure for arborescences, which is the reason why skeletons must obey this restriction.

There are many possible skeletons that can be defined for a set of body parts, but in practice we find that optimal results are obtained when the depth of the skeleton graph is kept low (to reduce inter-node dependencies during matching) and the lines formed between the nodes actually overlap with the animal’s morphology in the image (making curved body parts like rodent tails particularly challenging without intermediate keypoints).

### 3.4. Tracking

To address the **temporal association problem**, we devised a tracking algorithm that operates on grouped instances generated from the multi-animal pose estimation. The general algorithm is described in algorithm 1 which describes a standard multi-object tracking procedure. In brief, for each frame, we first generate a set of candidate instances from a window of recent frames that have been tracked, compute the matching cost between each candidate and each untracked instance in the current frame, perform the optimal matching and assign them to tracks.

To adapt this to the task of pose tracking specifically, we first employ one of two candidate generation functions: *flow shift* or *Kalman filter*. Inspired by Xiao et al. [33], the *flow shift* generator takes instances from previous frames and applies Farneback optical flow [14] to predict the displacement of the image between their respective frames and the current frame at the coordinates of the instance parts, thus generating a set of “shifted” instances with locations predicted by the image motion. This considerably improves the similarity between instances in the past and present ones, especially during bouts of fast social behaviors (e.g., chasing) during which the past location of one instance may more closely overlap with the current location of another. The *Kalman filter* generator uses instances tracked using the flow shift generator to initialize a filter model at the beginning of a track for each body part of each instance. At each subsequent frame, the Kalman filters are updated and candidate instances are generated by predicting the location of each past instance in the current frame. This motion model does not use image data, but can more accurately generate candidate instances when the appearance of animals change drastically. To compute the matching cost between instances, we use the *instance similarity* defined as

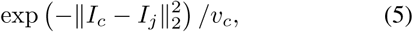

where *v*_*c*_ is the number of visible landmarks in the candidate instance.

#### Algorithm 1: Tracking Algorithm

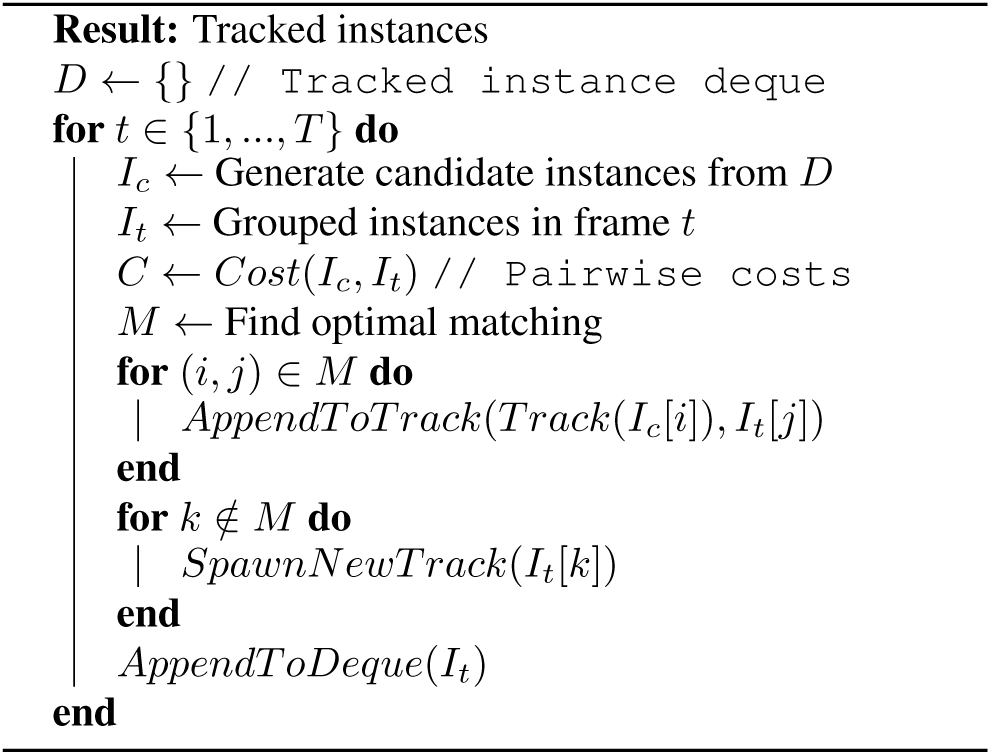

### 3.5. Neural network architectures

The neural network architectures we have implemented in our framework are agnostic to the approach employed. These *backbones* serve as generic feature extractors that can output feature maps, both CMs and PAFs, at any given resolution.

We employ the encoder-decoder meta-architecture (Figure 3a), a class of fully-convolutional network architectures that can describe most of the commonly used neural networks, such as those used in DeepLabCut [22] (Figure 3b) or UNet [29] (Figure 3c). We use this class of models as a means of defining a set of high-level hyperparameters that describe the *behavior* of the network rather than its explicit structure:

1. **Maximum receptive field size**: This is the largest spatial extent over which the network is theoretically capable of integrating features. This can be used to adjust the length scale over which the network should have the capacity to reason. The theoretical receptive field (RF) is increased by adding downsampling blocks in the encoder branch and can be calculated in closed form [4].
2. **Output resolution**: This determines the resolution of the output targets (such as confidence maps) which can determine the accuracy at inference time, particularly due to the quantization error inherent in subsampled outputs. This trades off with increased memory requirement, sometimes resulting in intractable training, particularly for bottom-up models. The output resolution is determined by the ratio of downsampling to upsampling blocks, where an equal number results in outputs at the same resolution as the input image.
3. **Representational capacity**: This determines the amount of parameters and compute that the network is able to use to capture image features across scales. It can be controlled by the base number of filters and by the filter rate, where a lower filter rate biases the distribution of representational capacity towards smaller features.

**Figure 3:**
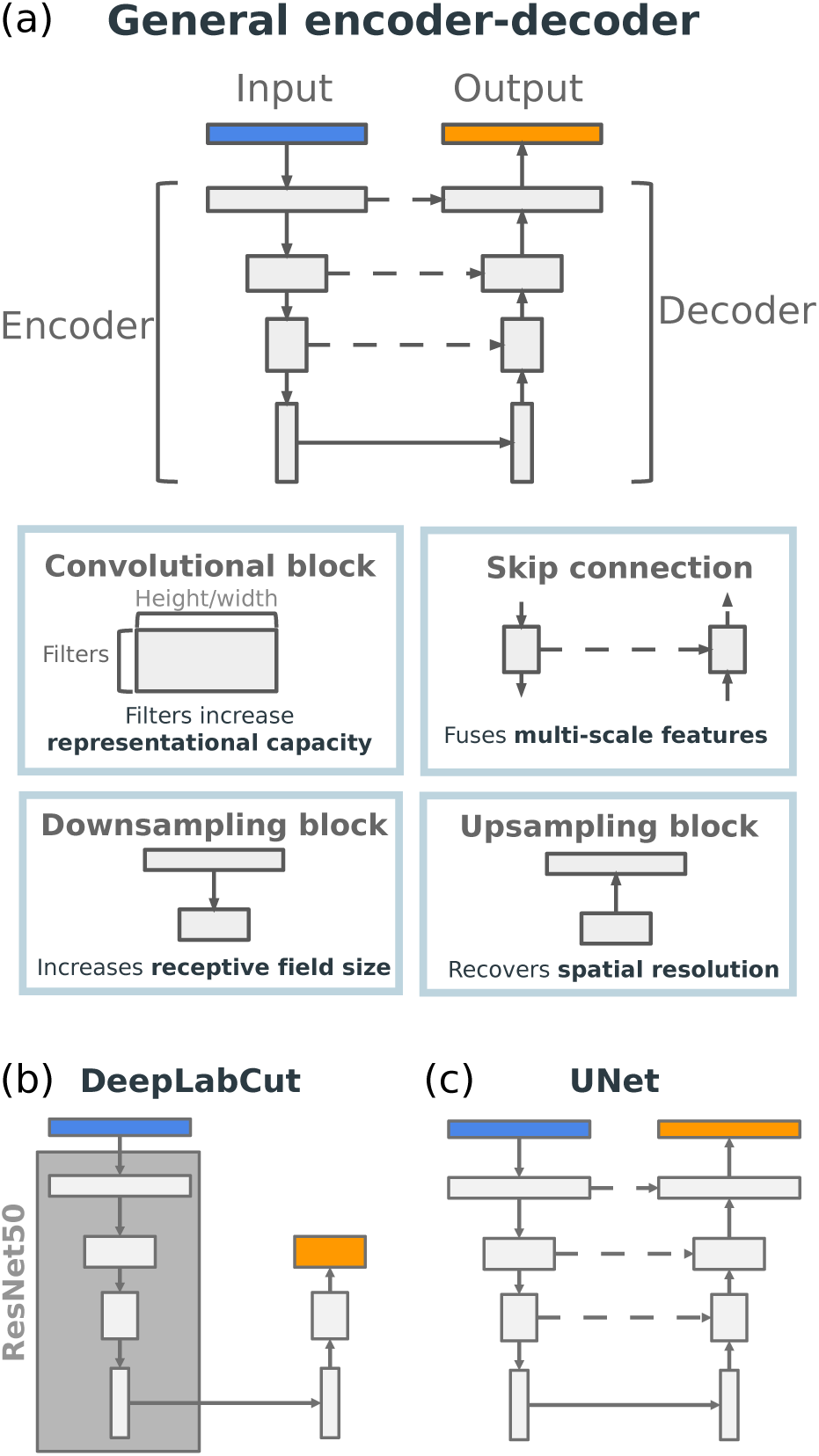
Neural network architectures generated by the encoder-decoder motif. **(a)** Encoder-decoder models are formed by a set of blocks that control specific properties of the architecture. These can be combined to achieve desired feature extraction properties, such as receptive field size or output resolution. **(b)** The standard DeepLabCut architecture. **(c)** The standard UNet architecture.

These hyperparameters make it simpler to select an appropriate architecture based on the properties of the data, rather than through careful engineering or black-box network architecture search methods.

We primarily use the generalized UNet as a base network [29] as it provides the flexibility to control these high level hyperparameters while implementing them with simple architecture design specifics, namely: *simple convolutional blocks* consisting of a stack of Conv-ReLU-Conv-ReLU-MaxPool layers, and skip connections to fuse features across scales in the decoder. We use bilinear upsampling, rather than transposed convolutions, as it has been previously shown to be effective and compute efficient [17].

For the ResNet implementation, we precisely replicate the architecture specifics employed in DeepLabCut [22] to ensure fair comparisons. Unlike previous attempts to replicate their architecture [17], we implement ResNet50, 101 and 152 with the exact layers necessary to make use of the standard ImageNet pre-trained weights, but retain DeepLabCut’s ability to control the encoder feature resolution through the use of dilated convolutions. This has been previously shown to be important for fully convolutional tasks that re-use the standard ResNet backbone[8] which in its standard configuration reduces feature spatial resolution by 32×. This makes it difficult to upsample the features in the decoder, especially without the use of skip connections. We also implement the decoder used by DeepLabCut, which is composed of 1 or 2 transposed convolutions with large kernel size that directly upsamples the encoder’s feature maps to the output target resolution, rather than through repeated upsampling blocks. In practice, we find that smaller upsampling blocks using bilinear interpolation yield better performance and training stability.

## 4. Results

### 4.1. Datasets

In order to evaluate our framework across a diversity of experimental conditions and animal morphologies, we generated three different datasets of multi-animal social behavior: flies, mice, and bees. These datasets were labeled using the SLEAP GUI and aided by the human-in-the-loop workflow (Figure 2).

The **flies** dataset consisted of 30 videos of pairs of male and female fruit flies (*D. melanogaster*) freely moving within a domed chamber and engaged in natural courtship behavior for up to 30 minutes, or until copulation. The videos were recorded from above at 150 FPS with a frame size of 1024 × 1024 × 1 in grayscale using infrared illumination, at a resolution of 30.3 pixels per mm. This resolution is close to the *minimum required* to be able to reliably capture individual leg tarsi as they span roughly ∼25 microns in diameter [30], equivalent to about 0.75 pixels at this resolution. We emphasize these limits to highlight that the findings described here regarding feature scales should be interpreted not just at the imaging parameters we selected, but are tied to the underlying scale of the biological features that these pose models are trained to detect.

We labeled 2000 frames with 2 flies visible in every image. The dataset was randomly split into 1600*/*200*/*200 frames for training/validation/testing respectively. The skeleton we selected consisted of 13 nodes spanning the anatomy of the fly: [*head, thorax, abdomen, wingL, wingR, forelegL*4, *forelegR*4, *midlegL*4, *midlegR*4, *hindlegL*4, *hindlegR*4, *eyeL, eyeR*]; and 12 edges: [(*thorax* → *head*), (*thorax* → *abdomen*), (*thorax* → *wingL*), (*thorax* → *wingR*), (*thorax* → *forelegL*4), (*thorax* → *forelegR*4), (*thorax* → *midlegL*4), (*thorax* → *midlegR*4), (*thorax* → *hindlegL*4), (*thorax* → *hindlegR*4), (*head* → *eyeL*), (*head* → *eyeR*)].

The **mice** dataset consisted of 30 videos of pairs of male and female white mice (*M. musculus*) freely interacting in a homecage environment with bedding to encourage naturalistic courtship behavior for ∼5 minutes. The videos were recorded from above at 40 FPS with a frame size of 1280 × 1024 × 1 in grayscale using infrared illumination, at a resolution of ∼1.9 pixels per mm. Although this resolution could be reduced, as the finest feature captured (the tail) occupies several pixels in these images, this resolution addresses other challenges inherent in this dataset, namely the low contrast due to low power IR illumination and white fur color of the animals against the bedding material.

For this dataset, we labeled 1474 frames with either 1 to 2 mice visible. The dataset was randomly split into 1178*/*148*/*148 frames for training/validation/testing respectively. The skeleton we selected consisted of 5 nodes spanning clearly visible anatomical landmarks: [*snout, earL, earR, tb*(*tailBase*), *tt*(*tailT ip*)]; and 4 edges: [(*snout* → *earL*), (*snout* → *earR*), (*snout* → *tb*), (*tb* → *tt*)]. We chose not to label the legs or paws since they were very intermittently visible from a single camera above.

The **bees** dataset consisted of 18 videos of pairs of female worker bumblebees (*Bombus impatiens*) freely interacting in a petri dish with hexagonal beeswax flooring for up to 30 minutes. The videos were recorded from above at 100 FPS with a frame size of 2048 × 1536 × 1 in grayscale, at a resolution of ∼14 pixels per mm. Similar to the fly dataset, this resolution is close to the *minimum required* in order to reliably capture the finer features of the bees such as their tarsi tips and antennae, both of which are frequently employed in social interactions [16]. This presented a distinct challenge as the large body size of the bees (∼200 − 400 pixels) together with the finest feature sizes (∼2 − 4 pixels) requires models to simultaneously capture features across a wide range of length scales.

For this dataset, we labeled 804 frames with 2 bees always visible (though often overlapping during interactions). The dataset was randomly split into 642*/*81*/*81 frames for training/validation/testing respectively. The skeleton we selected consisted of 21 nodes spanning the anatomy of the bees: [*thor, head, abdo, Lant*1, *Lant*2, *Rant*1, *Rant*2, *f Lleg*1, *f Lleg*2, *f Rleg*1, *f Rleg*2, *mLleg*1, *mLleg*2, *mRleg*1, *mRleg*2, *hLleg*1, *hLleg*2, *hRleg*1, *hRleg*2, *Lwing, Rwing*]; and 20 edges: [(*thor* → *head*), (*thor* → *abdo*), (*head* → *Lant*1), (*head* → *Rant*1), (*Lant*1 → *Lant*2), (*Rant*1 → *Rant*2), (*thor* → *f Lleg*1), (*f Lleg*1 → *f Lleg*2), (*thor* → *f Rleg*1), (*f Rleg*1 → *f Rleg*2), (*thor* → *mLleg*1), (*mLleg*1 → *mLleg*2), (*thor* → *mRleg*1), (*mRleg*1 → *mRleg*2), (*thor* → *hLleg*1), (*thor* → *hRleg*1), (*hLleg*1 → *hLleg*2), (*hRleg*1 → *hRleg*2), (*thor* → *Lwing*), (*thor* → *Rwing*)].

Together, these datasets comprise a range of imaging conditions and species morphologies which span a variety of image recognition dimensions. The **flies** dataset consists of small features (the inputs cannot be downsampled without considerable information loss), which occupy a small fraction of the field of view (160 × 160 bounding boxes out of 1024 × 1024 frames) resulting in a large number of irrelevant background pixels. The **mice** dataset, given our imaging conditions, consists of mostly coarse features that are difficult to precisely localize and occupy a relatively large proportion of the field of view with long-range dependencies between features (e.g., tail tip and rest of the body). The **bees** dataset is particularly challenging due to both fine and large scale image features, while occupying a relatively small portion of a very large field of view. Together, these conditions represent many of the common image recognition challenges inherent to social behavioral data collected in the lab.

These datasets and selected models optimal for each are available at sleap.ai.

### 4.2. Training procedure

All models were trained within the SLEAP framework at version 1.0.x (https://github.com/murthylab/sleap/releases/tag/v1.0.8) which uses Tensor-Flow 2 and Python 3.6. At the time of training we used TensorFlow 2.1, but all models are forward compatible with the current TensorFlow 2.3. We use the same base set of hyperparameters for all training runs. For optimization, we set the batch size to 4, define an epoch as a full iteration over the training dataset (with epoch boundary-respecting shuffling), and train for a maximum of 200 epochs with early stopping if the validation loss does not improve by at least 1 × 10^−6^ for 10 epochs. We use the Adam optimizer with AMSgrad enabled and an initial learning rate of 1 × 10^−3^ which we reduce by a factor of 0.5 after 5 epochs without a minimum validation loss decrease of 1 × 10^−6^ followed by a 3 epoch cooldown period during which the loss is not monitored. For consistency across all experiments, we apply only rotational augmentation (−180° −180°) to the data, though in practice we did observe that applying contrast-based augmentation improved the models’ ability to generalize to new data with slight lighting variations. Model checkpointing was triggered at the end of every epoch in which the validation loss improved by any amount and the final checkpoint was used for all experiments.

All training was performed on a single GPU, either locally with NVIDIA Titan RTX or on the cloud and our on-campus cluster with NVIDIA Tesla P100s. In both cases, GPU memory was typically not a limitation with our batch size, but we chose to keep it relatively small to ensure that training results could be reproduced on lower memory GPUs. The best performing models for each dataset were all able to be trained on Google Colab (which provides P100s) as well as locally on NVIDIA GeForce 1080 GTX Ti and 2080 RTX Ti cards with 8 GB of memory. System memory was not found to be a constraint and most of our training environments had 16 GB or less of available RAM.

### 4.3. Evaluation

In order to evaluate accuracy, we adapted several metrics that are standard in the human pose estimation literature. In particular, we adapt the Object Keypoint Similarity (OKS) metric [28] which is used nearly ubiquitously in human pose estimation benchmarks[21], but has not to our knowledge been previously adapted to animals. We compute the OKS in its standard form as described in Ronchi et al. [28]:

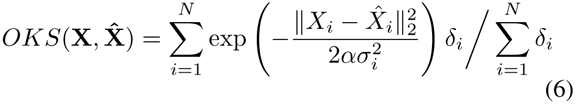

where **X** and 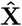 are the ground truth and predicted instance coordinates, respectively, for an instance with *N* nodes. The *δ*_*i*_ denotes the visibility, which is 0 if the node is missing from the ground truth instance. The inner term essentially expresses the distance from the ground truth coordinate as the posterior of a Gaussian with two scaling terms: *α*, the bounding box area occupied by the GT instance, and *σ*_*i*_, an empirically derived estimate of the labeling uncertainty for a landmark type *i*.

This last parameter, *σ*_*i*_, is intended to normalize the distance between prediction and ground truth with respect to the variance in localization consistency across human annotators, thus affording hard-to-label body parts such as the “left hip” in humans (which is subcutaneous and typically covered by clothing) greater leniency in the error calculation. The values of *σ*_*i*_ were originally derived from thousands of crowdsourced annotations for the standard 17 human keypoints in the MS-COCO dataset [28], however obtaining estimates for the human labeling variability for each new animal body part would be prohibitively laborious. Instead, we opt to set *σ* = 0.025 for all keypoint types, the standard deviation of human annotator uncertainty for the *easiest* keypoint: the left eye. We reasoned that this is a conservative value to use, where we essentially assume that all labeled animal body parts are as easy and consistently localizable to human annotators as the human left eye. This is certainly not the case for multiple body parts in our datasets (particularly in the **mice** dataset which has blobby features), so we consider this OKS as the lower bound of the true accuracy we obtain. The advantage is that this formulation of OKS can be interpreted to have comparable units (ranging from 0 to 1, where 1 is perfect accuracy) and variance scaling to those reported in the human literature, bridging the gap in evaluation metrics for the domain of animal pose estimation.

For evaluation of multi-instance pose estimation and tracking accuracy, we adopt the same procedures as employed in the widely used PoseTrack benchmark for human pose tracking [2]. Specifically, we compute the overall mean Average Precision (**mAP**) using the same procedure employed for the PoseTrack benchmark and widely reported in the human pose literature, a metric originally described in the Pascal VOC challenge [13]. Briefly, mAP computation involves classifying each pairing of (greedily matched) GT and predicted instance as a True Positive or False Positive by using the OKS as a cutoff at each of the following thresholds: {0.50, 0.55, 0.60, 0.65, 0.70, 0.75, 0.80, 0.85, 0.90, 0.95}. Precision at a single threshold is calculated as *TP/*(*TP* + *FP*), and recall as *TP/*(*TP* + *FN*). All predictions are sorted by their OKS and the cumulative TPs and FPs are computed for each predictions and recall and precision values from these partial TPs and FPs, i.e., for each pair of GT and predicted instance. Then, a set of 101 recall thresholds are defined with even spacing from 0 to 1, and the best precision value for samples that fall below each recall threshold is retrieved from the data, yielding 101 precision values. The Average Precision (**AP**) is computed by taking the mean over all 101 precision values, whereas the Average Recall (**AR**) is simply defined as the best recall at the current OKS threshold. This procedure is repeated for all 10 OKS thresholds and the final mean Average Precision (**mAP**) and mean Average Recall (**mAR**) are simply the average of the AP and AR over all thresholds. Although their calculation is non-trivial, the mAP and mAR provide balanced estimates of consistently reliable precision and recall performance across many certainty thresholds.

An alternative metric of localization accuracy that is often reported is the Percent Correct Keypoints (PCK) metric, which is simply the fraction of predicted keypoints that are closer than some threshold of Euclidean pixel distance. This is typically reported as “PCKh”” in which distances are normalized by the size of the person’s head to account for instance scale. As a more general metric, we instead report the mean PCK (**mPCK**) which is calculated by taking the average of the PCKs at a range of thresholds: {1, 2, 3, 4, 5, 6, 7, 8, 9, 10}. This provides a more generalizable metric that can be calculated in the same way for all datasets.

Finally, we report the 95th percentile of the Euclidean distance errors as a convenient metric of the expected pixel distance from GT for extreme outliers, i.e., the “worst case”.

All accuracy metrics reported here are computed on the held out test set of each dataset.

For the tracking accuracy evaluation, we employ the same MOT metrics as in PoseTrack [2] and compute them using the py-motmetrics framework [18]. In particular, the MOTA metric is computed as:

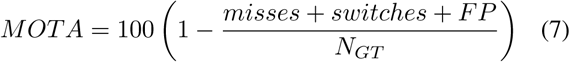

### 4.4. Multi-instance pose estimation approach

We conducted a series of experiments to explore the optimal approach for multi-instance pose estimation in each dataset and trained a total of 669 neural networks with varying approaches, network architecture hyperparameters, and training replicates. These results are summarized in Table 1 which lists the accuracy metrics for the best performing model for each dataset and approach. For the flies dataset, we find that the top-down approach works best, which we hypothesize is due to the requirement of fine feature descriptions in order to reliably capture the legs, benefiting little from the global context afforded by the bottom-up approach. For the mice dataset, the bottom-up approach considerably outperforms even the best top-down model. We hypothesize that this is due to the coarseness of the mouse features and long-range dependencies between body parts, in particular the tail, which strongly benefits from the global context encoded in the bottom-up approach. For the bees dataset, we see mixed results as expected due to the presence of both small and large scale features present in the dataset. While the overall accuracy as captured by the mAP and mAR metrics are much better for the bottom-up model which can provide increased context between body parts and across animals, the localization accuracy metrics mPCK and the 95th percentile of error distances reflect the benefit of the top-down model in emphasizing finer scale features over global structure.

**Table 1:** Best accuracy for each dataset and multi-instance approach. TD: Top-down, BU: Bottom-up.

| Dataset | Approach | mAP | mAR | mPCK | Error (95%) |
| --- | --- | --- | --- | --- | --- |
| Flies | TD | **0.832** | **0.881** | **0.878** | **2.78** |
| Flies | BU | 0.792 | 0.823 | 0.844 | 4.83 |
| Mice | TD | 0.401 | 0.527 | 0.571 | 27.0 |
| Mice | BU | **0.535** | **0.617** | **0.597** | **17.7** |
| Bees | TD | 0.658 | 0.759 | **0.589** | **14.6** |
| Bees | BU | **0.736** | **0.765** | 0.559 | 18.5 |

Overall, these results reinforce the premise that model accuracy is determined by dataset-specific characteristics and these should be considered when choosing a multi-instance approach.

### 4.5. Localization accuracy

Next, we sought to examine the performance of the best model for each dataset, to better understand how these dataset-wide summary metrics translate to the individual instance case. First, we visualized on example images from each dataset the distribution of errors in terms of Euclidean distance, by plotting circles with radii determined by the percentile of the error distribution in pixels (Figure 4a-c). We see that for the fly dataset, the vast majority of predictions fall within tight clusters around the ground truth location, suggesting that our error rates are likely to produce highly reliable estimates of the fly’s anatomy (Figure 4a). For the mouse dataset, we find highly accurate predictions for all body parts except the tail tip, whose extreme values fall within a much larger uncertainty radius, reflecting the difficulty in correctly predicting this body part (Figure 4b). For the bee, we see that some body parts are reasonably well-predicted, whereas others are not, particularly those centered on very fine scale features such as the mid-leg tips (Figure 4c) (note that the largest error circle reflects the 92.5th percentile rather than 95th for visualization clarity. We expect that these larger extremes may in part be due to the small size of the test set for this dataset).

**Figure 4:**
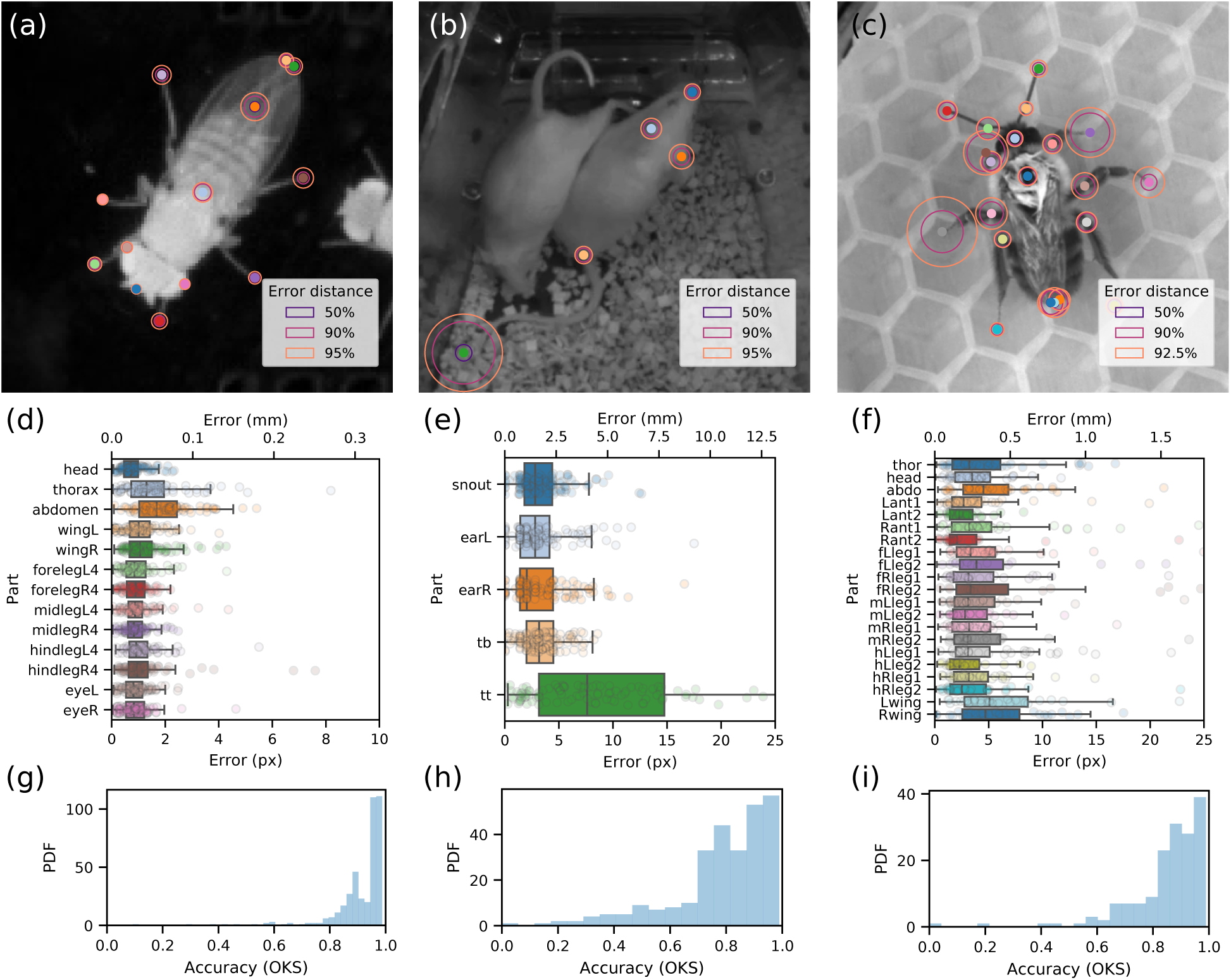
Multi-instance pose estimation accuracy. **(a-c)** Localization accuracy of best performing model on each dataset. Circle radii denote the percentiles of the Euclidean distance error distribution. **(d-f)** Distribution of part-wise localization error of best performing model on each dataset in both pixel and physical units. **(g-i)** Distribution of instance-wise accuracy as measured by the Object Keypoint Similarity (OKS) metric.

Plotting the distribution of the part-wise errors, we observe a more complete description of the first set of panels (Figure 4d-f). Note that the vast majority of points have error distances well below the anatomical scale of each animal, with nearly all of the points falling below 100 um for the flies (Figure 4d), 5 mm for the mice (Figure 4e) and 1 mm for the bees (Figure 4f).

Finally, the OKS score distributions for each dataset capture the variability in localization accuracy while accounting for the scale of the animals (Figure 4g-i). In particular, while more predictions fall into accuracy bins very close to 1.0 (the best possible score) than any other bin, the main determinant of the discrepancy in the mAP and mAR metrics appear to derive from the amount and spread of outliers—the long tail of the mice dataset (Figure 4h) accounts for the especially low mAP score. These distributions provide key insights into the source of errors for a given model, for example, by suggesting which kinds of instances or poses may benefit the most from additional labeled examples. For example, the bimodality of the fly OKS distribution (Figure 4g) points to two classes of errors — inspection of examples reveals that the lower distribution reflects cases in which the posterior landmarks are missing, a systematic source of error that arises from times when the male is closely interacting with the female such as during attempted copulation.

### 4.6. Receptive field size

Guided by the observation that the distribution of instance sizes varied by dataset (Figure 5a), we sought to explore how one of our encoder-decoder model hyperparameters, the *maximum receptive field size* (**RF**), would impact the performance of neural network architectures on different datasets. The RF determines the length scale over which the network is able to learn features; small RFs cannot incorporate global context, whereas large RFs may deemphasize smaller features (Figure 3). As described previously, we designed neural network architectures with a target RF by varying the number of *downsampling blocks* in the network and offset the loss in spatial resolution in the outputs with additional *upsampling blocks* (Figure 5b). Overlaying the RFs that we tested on examplars from each dataset illustrates the spatial extent of each animal or interacting animals that may be captured by features at different RFs (Figure 5c).

**Figure 5:**
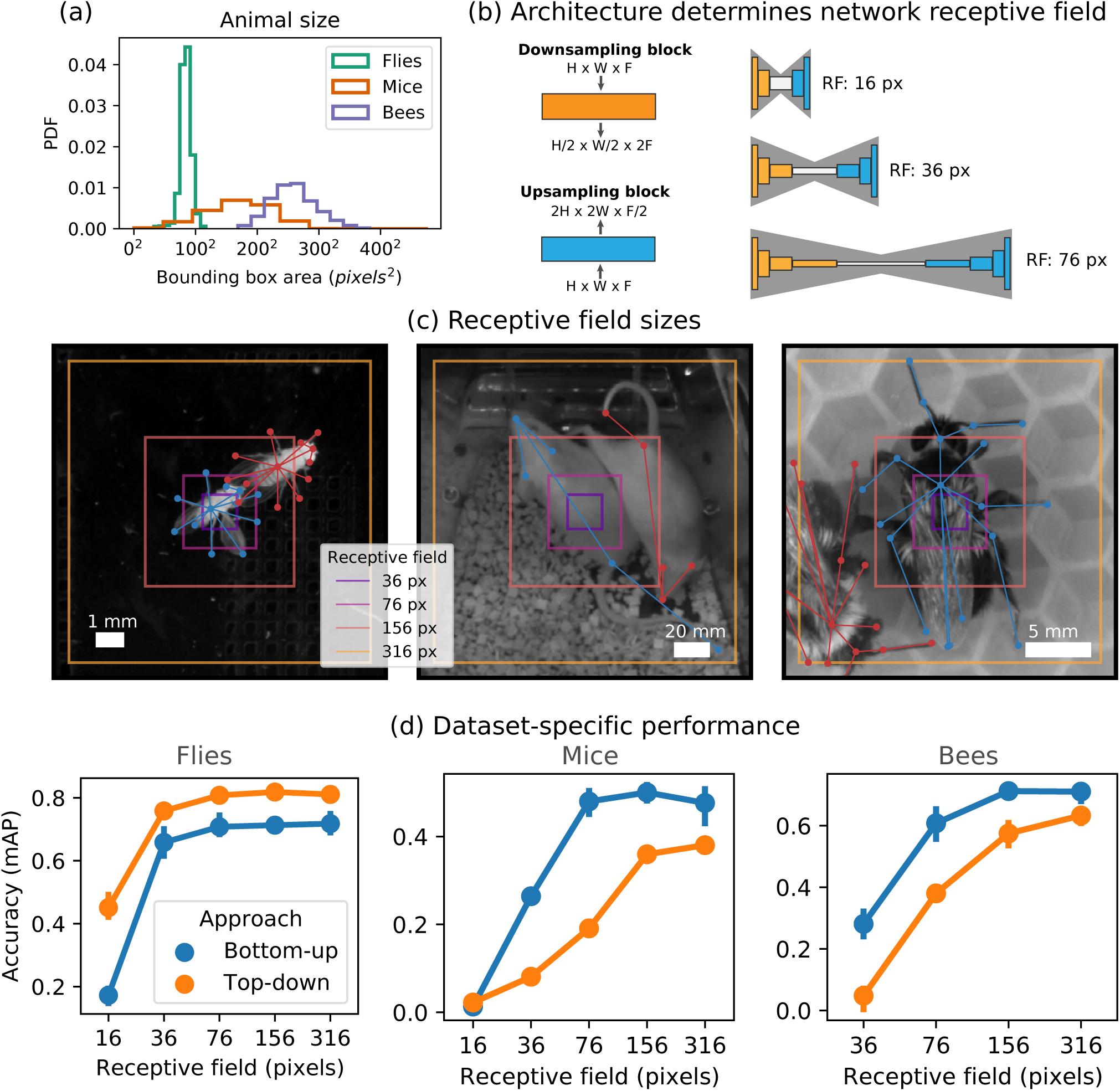
Optimal receptive field size and multi-instance approaches are dataset-specific. **(a)** Distributions of animal image area occupancy varies widely by dataset. **(b)** The maximum receptive field (RF) of a network sets a bound on the length scale of features that it can extract from images. Stacking simple blocks (*left*) enables the construction of network architectures with a desired receptive field size (*right*). **(c)** RFs visualized on an example from each dataset. While some animals have a small spatial extent (flies, *left*), others may require larger RFs to capture structure across animals (mice, *middle*) or even within the same animal (bees, *right*). **(d)** Accuracy of neural network architectures with different RF sizes vary by dataset and approach. Whereas flies benefit from top-down at small RFs, bottom-up networks with larger RFs perform better in mice and bees.

To test the effect of an architecture’s RF on each dataset, we trained models with varying maximum RF sizes and approaches for each dataset and report their accuracy as a function RF size (Figure 5d). We find that the optimal dataset-specific approach outperforms the alternative across all RF sizes, consistent with our hypothesis that the intrinsic properties of the representations induced by each approach are specifically suited to the characteristics of each dataset. As expected, all curves trend upwards as a function of increase maximum RF size. However, the diminishing gains in accuracy as RF increases suggests that there is an optimal RF size for integrating features in each dataset. Models with max RF size of 156 pixels appear to perform well across all datasets, suggesting that this may be a suitable common baseline architecture for general use. While the trend for top-down models for the flies dataset suggests that there is likely little to be gained by increasing the RF size – likely due to the small scale of the animals in that dataset (Figure 5a) – the trends for the other datasets indicate that increasing the RF may close the gap to the bottom-up model performances. This seems to be particularly salient for the bees dataset, which is expected due to its mixture of fine-scale and coarse features, the latter of which is perhaps not as easily captured in top-down models as it is in bottom-up models which explicitly encode global context.

### 4.7. Transfer learning

Previous work in single-animal pose estimation has proposed that transfer learning is critical for training large backbone networks, such as ResNet50, on few labeled examples, owing to the reuse of general purpose low level features (e.g., edges or image textures) [22, 23]. On the other hand, transfer learning may be dispensable for smaller, custom-designed networks [26]. To test how this applies to multi-animal pose detection, we also trained models with a ResNet50 backbone as the encoder, as well as employing strided convolutions to maintain feature map resolution, using randomly initialized weights or pretrained weights (i.e., transfer learning), and different approaches (in particular the bottom-up approach which has recently been implemented in DeepLabCut [22]).

We summarize our results by reporting the metrics for the best ResNet model in the pretrained or not-pretrained condition for each approach, as well as the best UNet-based model for reference (Table 2). These data indicate that pretraining can indeed improve final model performance, but these gains may be relatively small. For the flies dataset, the pretrained ResNet50 outperformed the randomly initialized version by 0.009 mAP and the optimal UNet by 0.011, but exhibited a higher 95th error distance percentile, suggesting that it fails to improve performance on the most difficult instances. For the mice dataset, sizable gains (+0.122 mAP) were observed in the top-down model when transfer learning was employed, but neither this model, nor the pretrained bottom-up ResNet model were able to match the optimal UNet (not-pretrained) model. For the bees dataset, gains were again observed with the use of transfer learning versus random initialization for ResNet50, particularly for the top-down model which seems to have greatly improved the localization accuracy — perhaps due to the ability of ResNets to efficiently preserve multi-scale features through their residual skip connections — but they still failed to outperform the optimal UNet model in terms of mAP, despite the overall improvement in localization accuracy.

**Table 2:** Performance of transfer learning for training multi-instance pose estimation models.

| Dataset | Approach | Backbone | Pretrained | mAP | mAR | mPCK | Error (95%) |
| --- | --- | --- | --- | --- | --- | --- | --- |
| Flies | TD | ResNet50 | **X** | 0.834 | 0.901 | 0.893 | 3.39 |
| Flies | TD | ResNet50 | ✓ | **0.843** | **0.912** | **0.900** | 3.07 |
| Flies | TD | UNet | **X** | 0.832 | 0.881 | 0.878 | **2.78** |
| Mice | TD | ResNet50 | **X** | 0.406 | 0.501 | 0.567 | 27.2 |
| Mice | TD | ResNet50 | ✓ | 0.528 | **0.621** | **0.611** | 22.7 |
| Mice | BU | ResNet50 | **X** | 0.253 | 0.324 | 0.388 | 59.0 |
| Mice | BU | ResNet50 | ✓ | 0.491 | 0.578 | 0.558 | 18.9 |
| Mice | BU | UNet | **X** | **0.535** | 0.617 | 0.597 | **17.7** |
| Bees | TD | ResNet50 | **X** | 0.650 | 0.744 | 0.518 | 17.3 |
| Bees | TD | ResNet50 | ✓ | 0.714 | **0.804** | **0.576** | **15.0** |
| Bees | BU | ResNet50 | ✓ | 0.651 | 0.682 | 0.417 | 18.6 |
| Bees | BU | UNet | **X** | **0.736** | 0.765 | 0.559 | 18.5 |

Altogether, these results indicate that while transfer learning may indeed provide improvements over random initialization for large network architectures (ResNet50), these networks still do not outperform smaller neural network architectures tuned to the properties of the dataset (Figure 5). Future improvements may involve the use of transfer learning together with optimal neural network architecture design.

### 4.8. Inference speed

Social behavioral monitoring data tend to suffer from two common performance bottlenecks: a large field of view (FOV) and high frame rates. Large FOVs are often required in order to permit animals to behave naturally, but this results in large image sizes (1024 × 1024 or larger) which are irreducible if the resolution is necessary in order to capture fine features of the animals, such as in the flies and bees datasets. High camera frame rates, in turn, are necessary since social behaviors can often occur on the order of milliseconds, resulting in huge videos, often on the order of > 100, 000 frames. This is different from human pose estimation, which uses small (*<* 512 × 512) images, often further sized down since most human body keypoints are still detectable at fairly low resolutions. On the temporal axis, “real-time” is often used to describe models that operate at > 30 FPS, the standard webcam frame rate, but models that operate at these rates would result in prohibitively slow inference times for animal behavior data. Both of these issues (large FOV and high frame rates) may lead to multiple-fold longer inference times than the experimental session itself, typically resulting in the need for additional compute infrastructure to deal with batch processing of experimental data.

Given these constraints, we sought to explore the range of possible inference speeds we could achieve with SLEAP. For these experiments, we benchmark inference times on top-down models trained on the flies dataset on a single desktop computer equipped with a NVIDIA Titan RTX (24 GB), 64 GB DDR4 RAM, Intel i7-6700K (4 cores), and Samsung SSD 950 PRO 512GB NVMe. To benchmark inference times, for each batch size and condition, we restart the benchmarking script which loads the models and performs an initial inference pass through a sample clip of 3200 frames to discount initialization time from TensorFlow’s autograph (this startup time is quickly amortized when performing inference on longer or multiple videos). Then, we perform 3 full inference passes through the data and store the mean runtime. We repeat this benchmarking procedure 3 times per batch size and condition to estimate the variability of multiple independent inference runs with separate initializations. The inference time includes the entire top-down inference pipeline, taking as input full resolution, raw video frames of size 1024 × 1024 × 1 and produces as output the coordinates of each body part for each instance in every frame of the video. We emphasize that the reported speeds constitute the complete *end-to-end* pipeline, which includes all preprocessing, forward pass of the anchor part detection model, instance anchored cropping, forward pass of the anchored-instance part confidence model, global peak finding, and refinement.

For the first experiment, we trained two sets of models: “accurate”” and “fast””. The former were trained with the highest accuracy (mAP = 0.832) UNet hyperparameters that we found during our previous experiments, whereas the latter had reduced filter sizes and output resolution resulting in reduced accuracy (mAP = 0.808), but require considerably fewer compute operations. We additionally benchmark the best ResNet50 top-down model which had comparable accuracy to the accurate UNet model (mAP = 0.835) (Figure 6a). We observe end-to-end inference speeds of 39*/*86*/*89 FPS for the ResNet/Accurate/Fast models, respectively, at batch size of 1. Inference greatly benefits from increased batch sizes as tensor operations can be optimally parallelized on the GPU at the cost of increased memory requirements. By increasing the batch size up to 128 images per batch, we observed speeds of 150*/*328*/*430 for the ResNet/Accurate/Fast models respectively.

**Figure 6:**
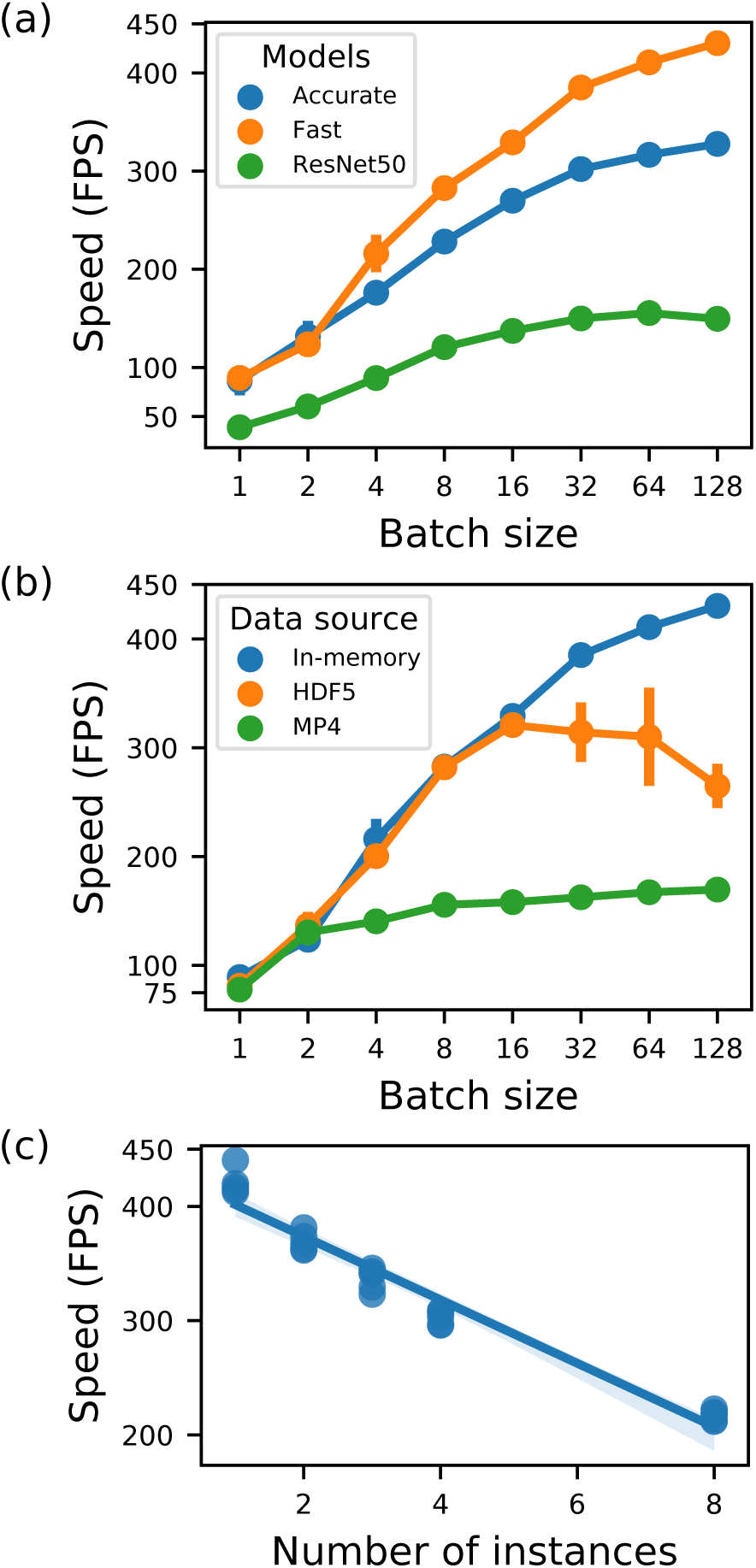
Multi-instance pose estimation speed. **(a)** Inference speed increases as a function of batch size and model configuration. The “fast”” models achieve 430 FPS at large batch sizes, at the cost of lower accuracy, whereas the “accurate”” models sacrifice speed (∼300 FPS) for greater accuracy. **(b)** Inference speed is bottlenecked by the source of the input data. When reading a high resolution MP4 video rather than performing inference on frames in memory, performance is limited by the video decoding speed to ∼150 FPS (*green*). At higher batch sizes, disk I/O becomes the bottleneck, resulting in the plateau of read speeds for uncompressed HDF5 frames. Inference speeds are measured with the “fast”” models from (a) at batch size 32. **(c)** Top-down performance scales with the number of animals in the frame. At batch size of 32 using the fast model, inference scales from 420 FPS with a single instance to 215 FPS with 8 instances.

Since these experiments were performed with the full video preloaded in memory, we next benchmarked the more realistic setting where I/O time is included. We benchmarked the fast model with three different data sources: memory, an uncompressed HDF5 dataset chunked by frame, and an MP4 file encoded with the `“superfast” libx264` preset (Figure 6b). The top speeds achieved when reading from the HDF5 dataset, which is read-pattern optimized due to the chunking, occurs at a batch size of 32 which resulted in 314 FPS as compared to the in memory speed of 385 FPS, reflecting the minimal overhead in reading from disk. Interestingly, decreased and more variable performance was observed at higher batch sizes, perhaps due to suboptimal disk access pattern for high bandwidth/frequency reads. Loading the image frames from the MP4 file reduced disk read overhead but was capped by the CPU decoding of the x264 compressed frames to a peak speed of 170 FPS at batch size 128. Perhaps further improvements could be obtained by moving the decoding to the GPU to minimize the CPU bottleneck.

Finally, since all our experiments used videos with two animals, we sought to characterize how the performance scaled with the number of instances in the frame (Figure 6c). We collected a new dataset of different numbers of flies interacting in the behavioral chamber. Previous top-down multi-instance pose estimation performance results did not include the instance cropping time and instead reported the performance in terms of crops per second [17]. Although it is theoretically expected that top-down inference speed will scale linearly with the number of animals, this has not been previously reported for a full *end-to-end* pipeline. We benchmarked the fast model at a batch size of 32 with videos of 1, 2, 3, 4 and 8 instances per frame and found that inference speed does indeed scale linearly with number of animals with a slope of −27.8 FPS per animal.

We note that although the machine used for these analyses is a high-end workstation, we have achieved comparable results with consumer-grade GPUs such as the NVIDIA GeForce 2080 RTX Ti. In addition, the upcoming generation of NVIDIA graphics cards that now come equipped with tensor cores provide huge boosts to neural network inference performance.

### 4.9. Tracking

The SLEAP tracking module employs two separate candidate instance generation approaches: *flow shifting*, which uses image information to predict the displacement of past instances, and *Kalman filtering*, which models the motions of the individual body parts in each instance to predict their location in the next timestep.

Here we report the tracking accuracy with the best approach (flow shifting or Kalman filtering) when applied to the flies and mice dataset (Table 3). For the flies, flow shifting is able to reliably predict the displacement of past instances, likely benefiting from the larger number of tracked body parts that are well defined (e.g., legs) as well as a much higher frame rate (150 vs 40). Since fly instances are easily trackable in the beginning of the session when they are less likely to interact (courtship interactions slowly ramp up over time), we evaluated the tracking accuracy only for the last 5000 frames (33 seconds) of the session, which ends at copulation and is typically the time when instances are most likely to be close. During this period, we find that the tracker commits 0.07 ± 0.69 identity switches per minute, a high variance likely resulting from a few particularly difficult to track examples. For the mice, we employ the Kalman filter and report the tracking accuracy on the full videos. We observe more switches per minute (1.26 ± 1.66), which is not intractable to proofread but leaves plenty of room for improvement.

**Table 3:** Tracking accuracy metrics.

| Dataset | Tracker | Videos | Switches<br>per min<br>(mean ±<br>sd) | MOTA |
| --- | --- | --- | --- | --- |
| Flies | Flow<br>shift | 95 | 0.07 ±<br>0.69 | 99.83 |
| Mice | Kalman<br>filter | 30 | 1.26 ±<br>1.66 | 99.98 |

## 5. Discussion

Here we have described SLEAP, a full-featured general-purpose multi-animal pose tracking framework designed for flexibility and tested on a diverse array of datasets representative of common social behavioral monitoring setups and on a wide range of organisms, from insects to vertebrates.

In future work, we intend to explore a broader class of backbone architectures, such as encoder-decoders with more sophisticated block types (e.g., residual or dense), as well as existing architectures shown to be effective at the task of human pose estimation such as HRNet [9] which may be well suited to address the challenges in multi-scale feature integration.

Although we demonstrated the capability of SLEAP to achieve end-to-end inference speeds compatible with realtime/closed-loop applications, we have not yet tested our models in such applications.

Extending SLEAP to tracking in 3D from multiple views may be another the direction of future work, though existing 3D pose tracking methods that build off of 2D predictions can already be configured take advantage of SLEAP [20].

Finally, a particularly important component to develop in future work with SLEAP will be to incorporate learnable tracking to enable the pose estimation models to better take advantage of temporal context. For example, the PAF representation could be extended to the time domain [27]. The top-down approach can also combine detection and tracking [31], although this requires sets of contiguous ground-truth frames which greatly increases the time and effort required for labeling.

## Acknowledgments

TDP is supported by NSF GRFP (DGE-1148900) and the Princeton Porter Ogden Jacobus Fellowship. ALF is funded by NIMH R00 MH109674, a Brain and Behavior Research Foundation award, and a Sloan Foundation award. ZYW is supported by the Princeton Catalyis Initiative, and SDK is supported by an NIH New Innovator DP2 GM137424-01 and NSF DEB 1754476. MM and JWS are supported by an NIH BRAIN Initiative R01 R01 NS104899, an NSF Physics Frontier Center grant (NSF PHY-1734030), and a Princeton IP Accelerator Award. MM is also supported by an HHMI Faculty Scholar award and an NIH NINDS R35 research program award.

We extend a special thanks to all of the SLEAP beta testers who graciously devoted time and effort to helping us develop the software framework.

## Notes

https://sleap.ai

### Competing Interest Statement

The authors have declared no competing interest.

